# A frontal cortical network is critical for language planning during spoken interaction

**DOI:** 10.1101/2023.08.26.554639

**Authors:** Gregg A. Castellucci, Christopher K. Kovach, Farhad Tabasi, David Christianson, Jeremy D.W. Greenlee, Michael A. Long

## Abstract

Many brain areas exhibit activity correlated with language planning^1–9^, but the impact of these dynamics on spoken interaction remains unclear. Here we use direct electrical stimulation to transiently perturb cortical function in neurosurgical patient-volunteers performing a question-answer task^10^. Stimulating structures involved in speech motor function evoked diverse articulatory deficits, while perturbations of caudal inferior and middle frontal gyri – which exhibit preparatory activity during conversational turn-taking – led to response errors. Perturbation of the same planning-related frontal regions slowed inter-speaker timing, while faster responses could result from stimulation of sites located in other areas. Taken together, these findings further indicate that caudal inferior and middle frontal gyri constitute a critical planning network essential for interactive language use^1^.

---

Turn-taking is a central feature of conversation across languages and cultures^11–14^. This key social behavior requires numerous sensorimotor and cognitive operations^11,15,16^ that can be organized into three general phases: comprehension of a partner’s turn, preparation of a speaker’s own turn, and execution of that turn. Using intracranial electrocorticography, we recently demonstrated that neural activity related to these phases is functionally distinct during both task-based and unconstrained turn-taking^1^. The networks active during the perceptual and articulatory stages of turn-taking consisted of structures known to be critical for speech-related sensory and motor function^17–26^, while a separate region of frontal cortex was active when participants planned their turns. Specifically, we found planning dynamics were most regularly observed in caudal inferior frontal gyrus (cIFG) and middle frontal gyrus (cMFG).

Given these observations, we hypothesize that cIFG and cMFG are necessary for language-related planning; therefore, perturbation of preparatory activity within these structures should result in behavioral deficits selectively related to planning spoken output. In animal models, disruption of planning dynamics in premotor regions can lead to slower^27^ and incorrect^28,29^ non-vocal responses that are otherwise executed properly. Likewise, perturbations of white matter tracts and cortical regions thought to be important for planning in humans can lead to slower and erroneous task-based speech^8,30–34^. We therefore predict that manipulation of cIFG and cMFG should result in similar deficits in the context of spoken interaction. Specifically, because these areas are persistently active early during planning^1^, we expect that disordering this activity would disturb high-level preparatory processes (e.g., formulating semantic or lexical structure)^35^ and thus lead to response errors but not articulatory dysfunction. Additionally, because efficient planning is central to achieving the rapid inter-speaker timing characteristic of conversational turn-taking^10,11^, we anticipate that perturbing this network should disrupt this coordination by protracting preparation times.

## Perturbation of neural activity during an interactive language task

To test our predictions, we used 50 Hz bipolar direct electrical stimulation (DES)^17–23,30,8,36–39^ to rapidly and reversibly perturb activity at 58 sites in 23 neurosurgical participants (Extended Data Tables 1, 2). We stimulated broadly across both cortical hemispheres (48 sites located in the left hemisphere) using both surface electrodes (n = 31) as well as intracerebral stereo-EEG depth electrodes (n = 27), enabling us to assess the behavioral results of disrupting our putative planning network (i.e., cIFG and cMFG; n = 22 sites) and a range of additional cortical loci (e.g., Fig. 1e). Participants performed the critical information (CI) question-answer task^10,40^, which was also used in our electrocorticography study to isolate planning-related neural dynamics during interaction. The CI paradigm relies on the same core processes as conversational turn-taking and specifically requires speech perception, articulation, and all processes related to planning a one-word answer (e.g., conceptual structure planning, retrieval of wordforms, phonological encoding)^4,41^ to occur in temporally defined epochs (Fig. 1a). At each stimulation site, participants answered a verbally presented battery of 28 to 143 questions (65 ± 21, mean ± SD) as quickly as possible while DES was randomly delivered on approximately half of the trials (49 ± 3%; range: 42% −60%) (see Methods). Stimulation intensity ranged from 7.5 to 24 V (15 ± 4 V) across sites, and DES was applied for 3.1 ± 1.0 s during stimulated trials – corresponding to 85 ± 18% of total trial duration and 91± 3% of planning period duration. Taken together, our perturbations should be capable of manipulating preparatory and sensorimotor processes related to spoken interaction.

**Figure 1.**
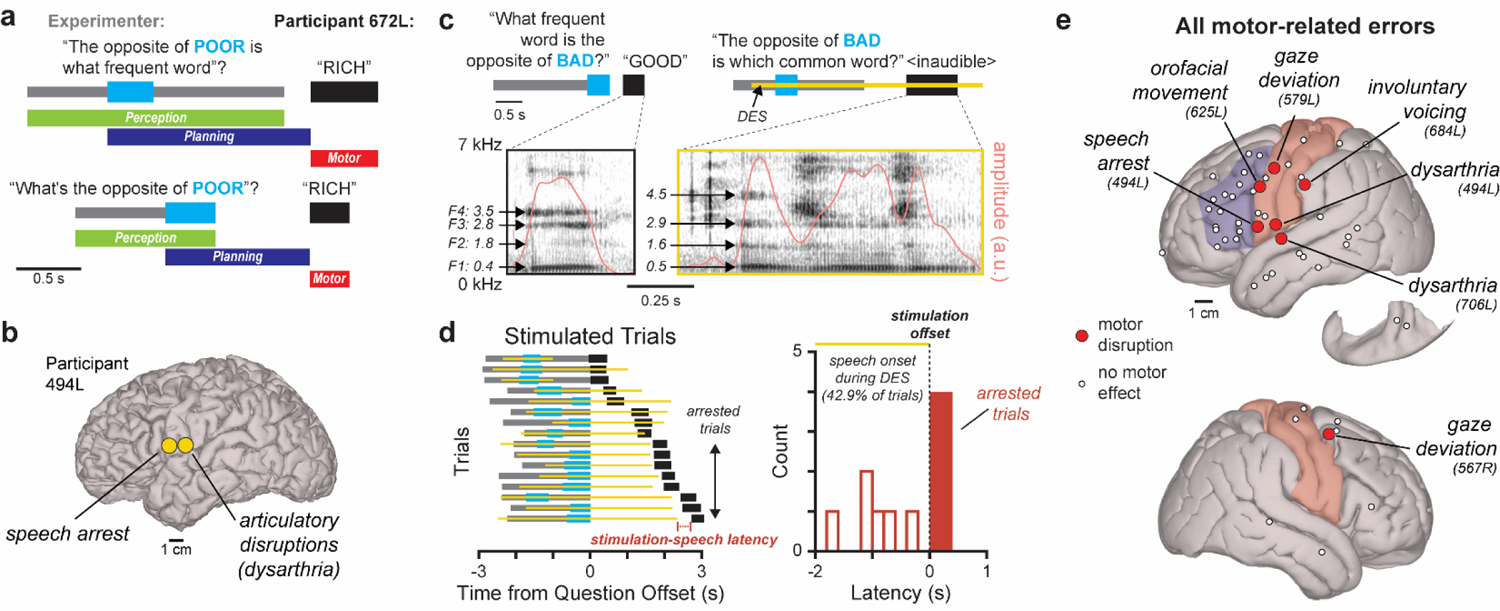
DES evokes motor deficits. (**a**) Example critical information (CI) questions where the CI is presented early (top) or late (bottom) with behavioral phases indicated. (**b**) Left lateral cortical surface of participant 494L with two stimulation sites indicated. (**c**) Example control trial (left) and trial where stimulation of the posterior site in (b) resulted in articulatory disruption (right); spectrograms of participant response with formant frequencies and envelope amplitude indicated (bottom). (**d**) Schematics of all trials where the anterior site in (b) was stimulated (left) with latencies between stimulation and speech onset shown at right. (**e**) Canonical cortical surface depicting all sites across participants where stimulation evoked motor deficits; all 6 stimulation sites within non-surface structures did not display motor disruptions (Extended Data Table 2). For (e), the location of precentral and postcentral gyri is approximated in light red, and the regions of caudal inferior and middle frontal gyri displaying neural activity related to language planning^1^ are approximated in light blue.

## Diverse motor deficits can result from DES

We first observed that cortical stimulation could elicit various motor disturbances that prevented performance of the CI task. For example, DES of two sites in participant 494L led to distinct deficits in articulatory performance (Fig. 1b). Specifically, stimulation of a site in subcentral gyrus, a region important for driving speech articulators^19,26^, resulted in unintelligible speech (i.e., dysarthria; Fig. 1c). Stimulation of another site in anterior precentral gyrus induced speech arrest, a well-documented DES-evoked motor deficit^19,22,23,39,42^ proposed to arise from an inability to initiate articulation^43^. In support of this interpretation, we found that answer onset occurred only 76 - 390 ms (191 ± 124 ms) following stimulation offset on arrested trials (Fig. 1d, Extended Data Fig. 1), which is faster than single word planning times^44,45^ and thus suggests that DES prevented the output of a pre-planned answer. We observed that stimulation evoked similar disruptions to overt movement – such as involuntary voicing and gaze deviations – in 5 other participants (Fig. 1e; Extended Data Table 2), and sites exhibiting such effects were preferentially clustered to areas important for orofacial motor function within precentral and postcentral gyri^17,22,39,46–48^ (actual: 86%; mean shuffled: 26 ± 16%) (p = 0.0007; permutation test). Our findings therefore indicate that these deficits resulted from selective perturbation of well-established motor networks^17–19,22,26^ that are distinct from regions performing other speech-related functions.

## DES can evoke planning-related errors

We next tested our first prediction that perturbing planning-related areas would elicit non-motor response errors, which are traditionally employed in DES studies as an indicator of disrupted language-related cognitive function^8,18,22,36,37,39,42,49,50^. We observed that DES often resulted in increased rates of semantic paraphasia (i.e., meaning-related errors; n = 11 sites) (Fig. 2a), anomia (failure to generate a spoken answer; n = 11 sites) (Fig. 2b), and hesitations (i.e., “umm”, disfluencies, filler words, see Methods; n = 22 sites) (Fig. 2c). Only a single site displayed an increased rate of neologisms (i.e., phonological errors) on stimulated trials (Extended Data Table 2), indicating that our perturbations largely affected high-level planning operations (e.g., formulating semantic and conceptual structure) as opposed to late-stage operations related to programming phonological and phonetic structure which immediately precede speech onset^9,35,41^. Likewise, stimulation-induced errors did not exhibit gross articulatory disruptions (e.g., Fig. 2a) and thus are unlikely to be related to motor-related dysfunction.

**Figure 2.**
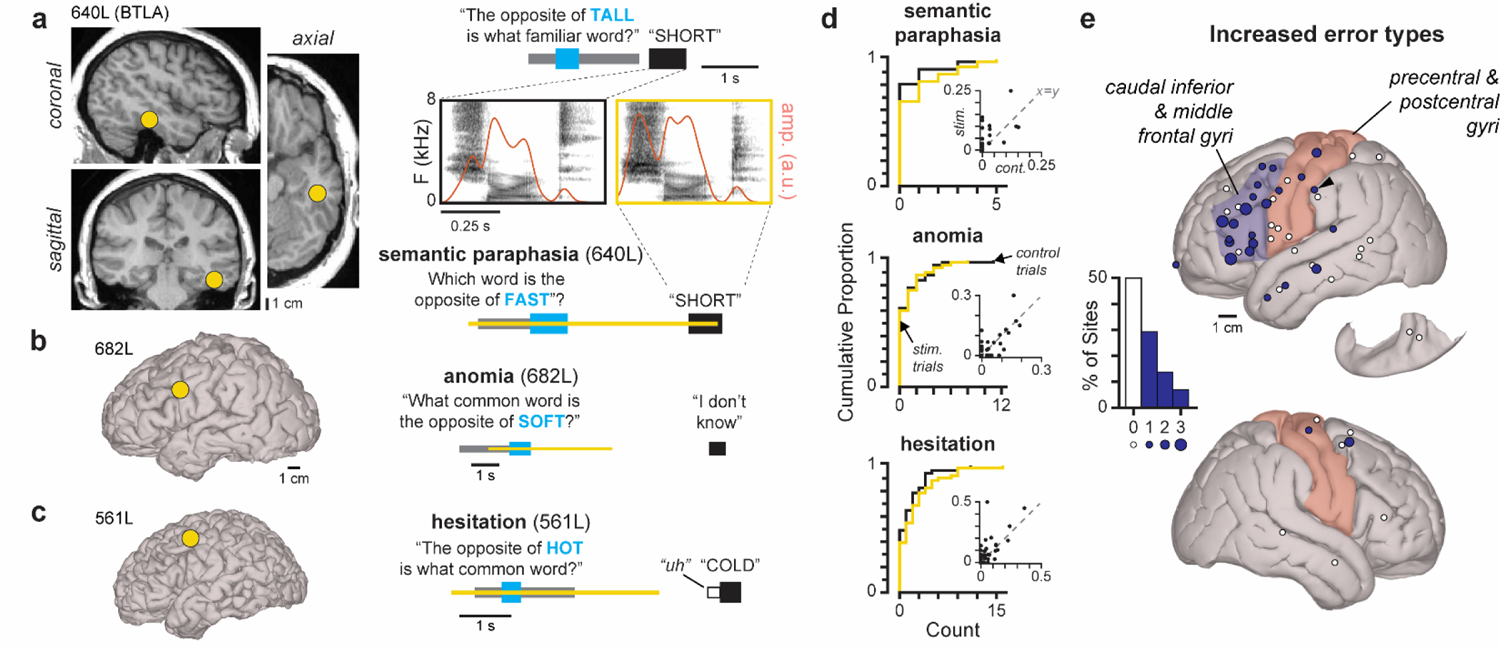
DES elicits planning-related response errors. (**a**) Preoperative magnetic resonance images depicting the stimulation site within the Basal Temporal Language Area (BTLA). At right, example control trial (top) and trial where BTLA stimulation resulted in semantic paraphasia (bottom); spectrograms and envelope amplitude of example responses are also presented (middle). (**b-c**) Example trials from sites where stimulation induced anomia (**b**) and hesitations (**c**); site locations indicated on participant cortical surfaces at left. (**d**) Cumulative distribution functions of error counts in control and stimulated trials across sites; inset scatterplots depict error rates. (**e**) The percentage of stimulation sites across participants displaying increased rates of one, two, or three error types (left), with corresponding locations depicted on canonical cortical surfaces (right); stimulation of sites in non-surface structures led to increases in either three error types (n = 1; BTLA site), one error type (n = 3), or no error types (n = 2) (Extended Data Table 2). Site displaying stimulation-induced neologisms is indicated with an arrow. For (e), the location of precentral and postcentral gyri is approximated in light red, and the regions of caudal inferior and middle frontal gyri displaying neural activity related to language planning^1^ are approximated in light blue.

In our interactive paradigm, we observed that errors were typically infrequent at individual sites, and their occurrence was variable across sites (Fig. 2d). Nevertheless, we found that DES significantly increased error rates within frontal and temporal regions (Fig. 2e, Extended Data Table 2), which is consistent with the findings of past neurosurgical studies that used non-interactive paradigms (e.g., picture naming, repetition)^18,22,39,42,51^. For example, stimulating the Basal Temporal Language Area (BTLA), an area of the fusiform gyrus thought to be critical for semantic access during speech planning^8,37,52,53^, resulted in semantic errors, anomia, and hesitations (Fig. 2a; Extended Data Table 2). Aside from this site, we found that 54% of loci where stimulation increased rates of any error type were located in cIFG (i.e., pars opercularis or pars triangularis) or cMFG, which is significantly more than expected by chance (mean shuffled: 39% ± 7%) (p = 0.0251; permutation test); likewise, 64% of sites displaying stimulation-induced increases in multiple error types were located within these regions. Therefore, while we found non-motor speech errors only sporadically resulted from DES, stimulation sites exhibiting such deficits were preferentially located in cIFG and cMFG, supporting our hypothesis that these regions are essential for proper language-related planning.

## Inter-speaker timing can be modulated by DES

We next sought to test our second prediction that perturbation of planning-related neural activity could protract the time required to prepare an answer during the CI task and thus disrupt inter-speaker temporal coordination. In particular, conversational turn-taking exhibits rapid transitions between speakers, with inter-turn gaps (i.e., floor transfer offsets) regularly 200 ms in duration or less^12,13^. Achieving these latencies requires speakers to simultaneously plan their responses while perceiving their partner’s speech^1,10,11^, a behavioral strategy that is also possible when CI is presented early in the question (e.g., Fig. 1a, top). During these early CI questions, participants typically respond to the experimenter with sub-second gaps that are significantly shorter than trials where the CI is presented late (Extended Data Fig. 2a), indicating that participants indeed began planning earlier when provided the opportunity. We therefore focused our gap timing analysis on early CI questions, and first tested how this metric was modulated by DES of an established speech planning site (i.e., BTLA). We found that stimulation of the site within BTLA (Fig. 2a) significantly lengthened gaps (Fig. 3a,b) (p < 0.0001, rank-sum test) without obviously affecting articulation (e.g., Figs. 2a and 3a), indicating that perturbation of regions important for language-related planning can measurably disrupt inter-speaker timing.

**Figure 3.**
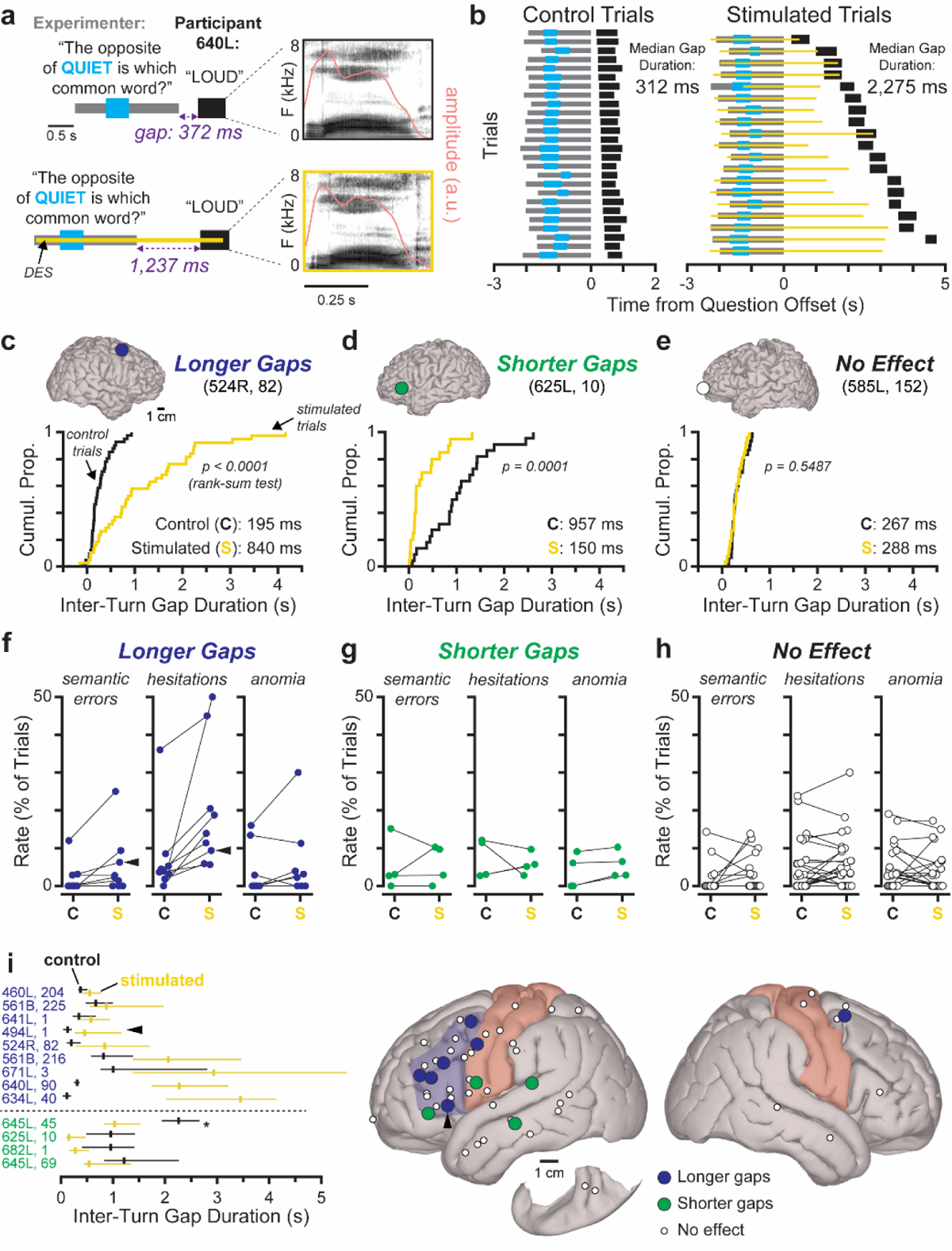
DES-induced modulations of inter-turn gap duration indicate a planning network. (**a**) Example control trial (top left) and trial where the Basal Temporal Language Area (BTLA; see Fig. 2a) was stimulated (bottom left) with inter-turn gap durations indicated; spectrograms of participant response with envelope amplitude overlaid (right). (**b**) All trials related to stimulation of the site in (a) sorted by gap duration. (**c-e**) Cumulative distribution functions of gap duration for example sites where stimulation induced longer gaps (**c**), shorter gaps (**d**), or no statistical difference in gap duration (**e**); site location indicated on participant cortical surfaces at top. (**f-h**) The error rates in control and stimulated trials for all sites where stimulation resulted in longer gaps (**f**), shorter gaps (**g**), or no effect on gap duration (**h**). (**i**) Median and interquartile range of gap duration for all sites where stimulation significantly affected this measure (left); site where DES induced shorter gaps on late CI trials only indicated with an asterisk. Participant and site number indicated in leftmost column (i.e., participant #, site #). At right, canonical cortical surfaces depicting the effect of stimulation on gap duration in all sites across participants; stimulation of sites in non-surface structures either lengthened gaps (n = 2; i.e., BTLA and posterior cingulate gyrus) or did not affect gap duration (n = 4) (Extended Data Table 2). The location of precentral and postcentral gyri is approximated in light red, and the regions of caudal inferior and middle frontal gyri displaying neural activity related to language planning^1^ are approximated in light blue. For (f) and (i), the site which displayed stimulation-induced perceptual deficits (see Extended Data Fig. 3c-g) is indicated with an arrow; note that this site was excluded from analyses of anomia in (f).

Across all sites, we observed that stimulation could result in significantly longer gaps (n = 9; Fig. 3c) or shorter gaps (n = 3 and an additional site in late CI trials only; Fig. 3d; Extended Data Table 2), with the remaining sites leading to no measurable changes (n = 38; Fig. 3e; Extended Data Fig. 2b). Because DES could increase or decrease gap duration, we tested whether either type of timing alteration was associated with planning-related response errors (e.g., Fig. 2a-c). We found that all locations where stimulation lengthened gap duration exhibited more anomia, semantic paraphasia, and/or hesitations on stimulated trials than control trials, with all such sites displaying an increase in at least one type of error (Extended Data Table 2) – significantly more than expected by chance (actual: 100%; mean shuffled: 48 ± 15%) (p = 0.0006; permutation test). Conversely, a consistent trend for increased errors was not observed when stimulation resulted in shorter gaps (75%, p = 0.2795; permutation test) or no effect in gap duration (42%, p = 0.9412; permutation test) (Extended Data Table 2). Therefore, perturbation of language-related planning via DES is likely signaled by a combination of longer inter-turn gaps and qualitative behavioral deficits including paraphasia, hesitations, and anomia.

We then examined the anatomical distribution of stimulation sites across participants to test whether DES-induced modulations in gap duration were spatially organized. We found that sites where stimulation did not affect inter-turn gaps were located broadly across the left and right hemispheres; in contrast, sites exhibiting stimulation-induced increases in gap duration were primarily restricted to frontal cortical areas (Fig. 3i). Excluding the site in BTLA (Fig. 2a), 7 of these 8 putative planning sites were located either in cIFG or cMFG – significantly more than expected if these sites were randomly distributed (actual: 88%; mean shuffled: 39 ± 16%) (p = 0.004; permutation test) – while a single site was located in left posterior cingulate gyrus (Extended Data Fig. 3a; Extended Data Table 2). Likewise, 32% of sites in cIFG and cMFG displayed stimulation-induced increases in gap duration while only 3% of sites outside these regions exhibited this behavioral deficit (shuffled mean: 14%). These results therefore confirm our second prediction that activity within cIFG and cMFG is critical for maintaining rapid inter-speaker coordination. Meanwhile, sites where DES resulted in shorter gaps (Extended Data Fig. 3b) were located outside cIFG and cMFG and not clustered to any specific region of cortex (Fig. 2i), suggesting this phenomenon does not arise from the same mechanism underlying stimulation-induced gap lengthening.

Finally, because we are the first to examine the effects of DES on inter-turn gaps, we confirmed that the observed increases in gap duration were unlikely to have arisen from factors other than perturbed planning. For example, participants in this study were undergoing surgical treatment for either epilepsy, brain tumors, Parkinson’s disease, or essential tremor, and the constraints for these procedures required DES to be delivered using three different methods (Extended Data Table 1). However, we found no significant relationship between the probability of observing stimulation-induced gap lengthening and the clinical condition of the participant (χ^2^ vs. constant model: 0.965, p = 0.617), the type of stimulator used (χ^2^ vs. constant model: 0.468, p = 0.791), or stimulation intensity (χ^2^ vs. constant model: 1.370, p = 0.242), indicating that this putative planning deficit did not arise from these unconstrained factors. Increased inter-turn gap durations were also unlikely to have resulted from disrupted language comprehension, as perceptual deficits were observed at only a single stimulation site examined in this study (Extended Data Fig. 3c-e). Notably, DES of this particular locus in pars triangularis also evoked a combination of longer gaps, semantic paraphasia, and hesitations (arrows in Fig. 3f,i; Extended Data Fig. 3f-g), indicating a mixed function related to perception and planning. This finding is in agreement with our previous intracranial recordings, which identified a subregion of cIFG overlapping with ventral pars triangularis that was active during the preparatory and perceptual phases of the CI task^1^.

## Discussion

In summary, we confirm our predictions that perturbation of cIFG and cMFG would 1) evoke non-articulatory response errors and 2) result in significantly longer inter-speaker gaps in an interactive speech task. Our findings therefore demonstrate that these regions are essential to planning during spoken communication and provide further evidence for the existence of a spatially and functionally distinct language-related planning network^1^. Previous studies have reported that DES of left cIFG (i.e., classical Broca’s region^54,55^) can result in a range of deficits related to cognitive and motor function during speech production^18,22,39,42,43,56^, and activity within this region has been linked to multiple linguistic and domain-general cognitive processes^3,5,9,54,57–63^. Therefore, while cIFG likely represents a heterogenous, potentially multimodal structure^60,63^, our results indicate this region contains circuitry essential for planning spoken language^1^. Similarly, cMFG stimulation has been observed to result in speech arrest and various non-motor errors^18,51,62^, but these behavioral effects often occur with less regularity when compared to DES of other frontotemporal structures^18,22,39,42^. Although regions of cMFG are known to be important for several non-linguistic functions^64–68^ including generating eye movements^46,47^ (e.g., Fig. 1e), its role in language has remained unclear. However, a recent large-scale imaging study found that a subregion including cMFG and adjacent precentral gyrus (i.e., Area 55b) displays activity during linguistic tasks and exhibits strong connectivity to language-related circuitry^69^. Likewise, cMFG resection can lead to transient^70^ or long-lasting^71^ deficits in verbal fluency, suggesting this area may contribute to speech production. Our stimulation and intracranial recordings further these important findings by demonstrating that cMFG is critically involved in the planning of spoken language^1^. While additional research is required to determine the exact planning operations^2,35,72^ performed in cMFG and to define the boundaries of its language-related territory, its close proximity to speech areas in cIFG^69,73^, dorsal laryngeal motor cortex^20,38^, and middle precentral gyrus^9^ further motivate this region as a key node within the human language network.

Our broad sampling of the brain with DES and targeted measurement of inter-speaker timing revealed four sites where cortical stimulation resulted in significantly faster participant responses (Extended Data Fig. 3b). Previous studies have reported that perturbation of cortical structures can improve or speed up task behavior in humans and animal models, suggesting that this phenomenon is generalizable across species and behaviors^74,75^. For example, speech rate could either increase or decrease when language-related premotor regions were focally cooled^76^. We found that several sites outside of regions related to action and speech elicited faster responses; therefore, disrupting neural activity via DES may have modulated a variety of distinct processes both related and unrelated to language – such as global arousal, impulsivity, planning, and/or language comprehension – which ultimately led to faster responses. While the underlying mechanisms remain unknown, these data provide promising initial evidence for cortical stimulation as a therapy for augmenting cognitive function, perhaps in a manner analogous to the usage of deep brain stimulation and transcranial magnetic stimulation to treat motor and nonmotor symptoms in a range of disorders^77–79^.

In conclusion, this study advances our understanding of the human language network by establishing cMFG as a critical speech-related planning area and confirming that cIFG causally contributes to the preparation of spoken language. Our results suggest these regions contribute to generating the meaning of speech (e.g., conceptual or lexical content) rather than its lower-level structure (e.g., phonological or phonetic content), as perturbing these regions elicited both slower responses and high-level errors (i.e., semantic paraphasia, hesitations, anomia) but not neologisms or articulatory errors. Therefore, cIFG and cMFG likely function as a key module in a larger network of both language-specific circuitry, such as those involved in encoding phonetic and phonological form^2,4,7,9,41,80–82^ as well as areas involved in general task performance^5,60,61^ – for example, the longer inter-turn gaps resulting from posterior cingulate stimulation (Extended Data Fig. 3a) may have resulted from disrupted cognitive control^83,84^. Further DES research combined with sensitive, ethologically relevant behavioral metrics is well-suited to identify additional components of this planning network and to delineate their precise contributions to typical and disordered language generation^85–87^.

## METHODS

### Participants

Study participants were patient-volunteers undergoing surgical treatment at the University of Iowa Hospitals and Clinics for medically intractable epilepsy, brain tumors, or implantation of deep brain stimulation (DBS) electrodes. Data from participants with previous brain resections or infarcts were not included in this study. During the course of treatment, neural activity was recorded using electrocorticography (ECoG) electrodes and/or intracerebral stereo-EEG depth (sEEG) electrodes either chronically (i.e., seizure focus determination) or acutely (i.e., awake craniotomy for tumor removal/epilepsy treatment or during a deep-brain stimulation electrode implant procedure). In ten participants, language function was confirmed to be either left lateralized or bihemispherically distributed using Wada testing or functional magnetic resonance imaging; the remaining participants were right-handed (n = 11) or left-handed (n = 2) with unknown lateralization for language. Participants consented to research, and the University of Iowa Institutional Review Board approved all procedures. Finally, all participants were native speakers of English except for one (684L), who was a native speaker of Vietnamese; consequently, this participant was only screened for stimulation-induced sensorimotor effects and their behavioral data were not analyzed further. Additional demographic and clinical information is available in Extended Data Table 1.

### Data Acquisition

For awake craniotomy patients, electrical signals resulting from direct electrical stimulation (DES) were recorded using subdural ECoG grids and/or strips manufactured by Ad-Tech Medical or PMT. Signals were amplified and sampled at 2034.5 Hz using a multichannel amplifier and digital acquisition system (PZ2 or PZ5 preamplifier with an RZ2 processer; Tucker-Davis Technologies). For chronically implanted epilepsy patients, electrical signals from subdural electrode grids, strips, or sEEG intracerebral depth electrodes (Ad-Tech) were recorded at 2000 Hz with a multichannel amplifier and digital acquisition system (Atlas system, Neuralynx). In both contexts, analog input channels synchronized with the neural recordings additionally acquired the output of 1-3 microphone(s) which captured the speech acoustics of the experimenter and participant. Input channels were typically sampled at 48,828 Hz by the TDT system and 16,000 Hz by the Neuralynx system and downsampled offline. In addition to the electrical signals, a video of the participant was also acquired at 24 fps for 20 of 23 experiments. The video was synced to the electrophysiological data after the experiment and provided a secondary high-quality audio recording channel, which was sampled at 48 kHz.

In a single awake craniotomy experiment, electrical signals were not recorded due to a recording system error and consequently only a video of the experiment was recorded (641L; see Extended Data Table 1). This video provided images of the stimulator and captured experiment acoustics; thus, stimulation and behavioral timing was estimated using this data stream.

### Direct Electrical Stimulation (DES)

DES was performed using a constant voltage stimulator (SD9, Grass Instruments). Charge balanced, biphasic pulses (0.2 ms duration) at 50 Hz were applied to the brain using either: 1) a handheld stimulator (Magstim BiPolar Probe; MVAP Medical Supplies, Inc., Thousand Oaks, CA) in awake craniotomy for tumor resection, 2) subdural ECoG strip electrodes (Ad-Tech) in awake craniotomies for deep brain stimulator implantation, or 3) subdural ECoG grid/strip electrodes or sEEG electrodes (Ad-Tech) in chronic epilepsy monitoring procedures. Because the position of the handheld stimulator could vary slightly from trial-to-trial (e.g., the stimulator could be removed momentarily to apply saline to the brain), we considered stimulation locations that varied by less than a centimeter to represent a single stimulation site. To titrate the stimulation voltage level, DES was first applied starting at ∼5V and increased in ∼2.5V steps until the participant reported a sensation, after-discharges were observed, or a maximum of 20-25V was reached. The highest voltage not resulting in a sensation or after-discharges was used as the stimulation level for the experiment. Occasionally, the stimulation voltage would be lowered during the course of an experiment if delayed after-discharges were observed. The stimulation voltage at each site included for analysis in this study is reported in Extended Data Table 2; in cases where the voltage was lowered, the lowest intensity is reported.

Stimulation timing was manually controlled by the experimenter, who also presented the questions to the participant in most cases – however, in some experiments, a separate experimenters presented questions and delivered stimulation. The experimenter controlling the stimulator typically began applying DES prior to CI presentation and stopped when the participant spoke their response. However, considerable variability in stimulation timing was observed across trials, and any trial where DES was applied between CI offset and answer onset was considered a ‘stimulated’ trial. In all cases, participants were not given any sensory cues related to the timing of stimulation (e.g., visual or auditory).

### Anatomical Reconstructions

Cortical surface reconstruction was performed using T1-weighted or magnetization-prepared rapid gradient-echo (MPRAGE) images obtained during the clinical workup and the ‘recon-all’ pipeline in FreeSurfer^1^. In cases of poor T1 imaging, FreeSurfer processing was repeated with combined T1 and T2 images or T1 images with fluid-attenuated inversion recovery (FLAIR) sequences to improve pial surface parcellation. White-matter segmentation errors resulting from low image quality or tissue abnormalities were corrected manually.

### Stimulation site localization in awake craniotomies for tumor resection

In one intraoperative tumor resection case (494L), stimulation site localization and coregistration was performed using intraoperative photographs, which were aligned to reconstructed cortical surface meshes via visual comparison of gyral anatomy by two raters (G.A.C. and J.D.W.G.). In all other cases, intraoperative photographs along with pre- and post-operative magnetic resonance (MR) images were used to localize both leads of the hand-held stimulator probe. Specifically, we first localized the craniotomy by 1) aligning intraoperative photographs to reconstructed cortical surfaces via visual inspection, 2) coregistering any burr holes and bone flap margins visible on the postoperative MR images to preoperative images using the FLIRT tool in the FSL package^2^, and 3) recording the coordinates from the Stealth neuro-navigation system (Medtronic, Minneapolis, MN, USA) used for craniotomy planning. Stimulation position was then defined on cortical surface renderings via visual comparison of gyral anatomy by two independent raters (F.T. and C.K.K) and cross-verification with a third rater (J.D.W.G.).

### Electrode localization in awake craniotomies for deep brain stimulator implantations

Localization of subdural ECoG strip electrodes (Ad-Tech, Racine, WI, USA) in patients undergoing deep brain stimulation (DBS) surgery was performed using a combination of intraoperative fluoroscopy, preoperative MR images, and pre- and post-implantation computed tomography (CT) collected for clinical reasons. Because the ECoG strips were temporarily placed during surgery and removed before closure, intraoperative fluoroscopy was required to record strip locations for surface-based localization while CT and MR images were used to reconstruct cortical surfaces and align the fluoroscopic images. First, all MR and CT images were converted from DICOM (Digital Imaging and Communications in Medicine) to NIfTI (Neuroimaging Informatics Technology Initiative) format using dsm2niix tool in MRIcroGL^3^. Preoperative CT scans were then employed as a 3D frame for the coregistration of 2D fluoroscopic images. Specifically, the ‘isosurface’ function in MATLAB was used to compute the surface geometry of the preoperative CT, and cortical surface meshes were then transformed into CT space. To coregister the fluoroscopic image, multiple common control points were identified in the CT volume and 2D fluoroscopic projection, including skull contours, the superior and inferior terminations of the frontal sinuses, the glabella, and screws of the stereotactic frame (CRW; Integra, Inc., Princeton, NJ). Because the view plane angle and field of view varied between fluorograms, control points were individualized based on the visible skull anatomy. For each individual, a minimum of four different control points were utilized. For cases with an insufficient number of plainly visible skull landmarks, a post-implantation CT which displayed DBS leads and burr hole covers was coregistered with the pre-implantation CT to provide an additional set of reference points. Next, an orthographic projection from the CT image space to the imaging plane of the fluoroscope was computed through constrained minimization of squared error between control points. This procedure yielded an orthogonal affine projection relating control points in CT space to those in the fluorograms. After this alignment, ECoG electrode locations were defined as the point at which a line orthogonal to the fluoroscopy-aligned view plane, passing through the electrode shadow visible in the fluorogram, intersected the cortical surface. Finally, the RAS coordinates of each electrode were transferred to Montreal Neurological Institute (MNI) space.

### Electrode localization in chronically implanted patients

Electrode localization in epilepsy patients undergoing chronic electrode implantation was performed by identifying characteristic metallic-induced susceptibility artifacts and punctuate radiodensities in post-implantation MR and CT images, respectively. Electrode coordinates were then transferred to pre-implantation images via linear image coregistration, followed by manually guided thin-plate spline warping to account for nonlinear imaging and structural distortions. Control points for warping were determined by visually identifying corresponding landmark coordinates in pre- and post-implantation imaging.

### Stimulation site coregistration

All contact locations were first determined on cortical surface renderings in Right-Anterior-Superior (RAS) coordinate space. Anatomical labeling of the electrodes was performed through surface-based coregistration and segmentation for each participant using the Desikan-Killiany-Tourville (DKT) atlas^4,5^ as assigned by FreeSurfer^6^. After automatic parcellation, electrode locations relative to DKT atlas labels were visually inspected by two raters (G.A.C and J.D.W.G) and corrected if necessary. To provide a consistent operational boundary between rostral and caudal middle frontal gyrus in line with our previous research^7^, raters considered any stimulation sites posterior to the extension of the anterior horizontal ramus of the inferior frontal gyrus as within caudal middle frontal gyrus and those anterior to this boundary to be within rostral middle frontal gyrus. Finally, preoperative T1 images were non-linearly coregistered to an MNI-aligned template brain (CIT168 template)^8^ using symmetric diffeomorphic registration in the ANTs toolbox^9^ to transfer electrode locations to MNI space.

Stimulation sites on the lateral cortical surface and superior temporal plane in individual participants were rendered on coregistration plots (e.g., Fig. 1e) by plotting the center of the two stimulation electrodes on the gyral surface of the MNI152 reference brain. The locations of sEEG stimulation sites were rendered on canonical and individual cortical surfaces by plotting the coregistered coordinates of the most superficial contact of the electrode shaft on the gyral surface of the MNI152 brain or the location where the shaft penetrated the gyral surface, respectively. SEEG stimulation sites located on the medial cortical surface, within insular cortex, or on the ventral cortical surface (i.e., “non-surface sites”; n = 6) were not rendered on coregistration plots. The coordinates for each pair of stimulated electrodes are reported in Extended Data Table 2.

### Behavioral Task

Participants completed the Critical Information (CI) task, which required them to answer simple questions as quickly as possible. The CI questions were read by the experimenter and adapted from a stimulus set used in our previous work^7^ that itself was adapted from an established Dutch stimulus set^10,11^. Each CI question contains a single word (i.e., the critical information) which is required to answer the question (Fig. 1a). In most experiments, questions required participants to generate the antonym of common words; however, in a subset of experiments, questions could also ask about animal sounds or body parts.

Questions were presented either in randomized order or in pseudorandomized order to avoid repeating the same CI on subsequent trials. On approximately half of the trials, DES was applied. In most experiments, stimulated and control (i.e., non-stimulated) trials were balanced such that the same question would be presented twice – once as a control trial and once as a stimulated trial. However, because of clinical considerations (e.g., time constraints in the operating room or to avoid after-discharges), question order, stimulation status, and wording often deviated from the planned protocol. In addition, most participants completed a block of the CI questions prior to the stimulation experiment to familiarize them with the task; these data were not analyzed in the present study. In one participant (460L), other tasks (e.g., repeating syllables, pressing a button) were interleaved with trials of the CI task.

The audio acquired with the electrophysiological acquisition system and/or video camera was annotated and timestamped by a trained phonetician (G.A.C.) to determine the onsets and offsets of all experimenter questions, CI, and participant responses. All final timestamping was performed blind in regard to stimulation timing. Questions which could not be accurately timestamped due to background noise (e.g., clinical team talking in background, operating room door closing, medical instrument alerts) were excluded from analysis. Likewise, any antonym question where the participant gave an answer using “not” and the CI (e.g., Question: The opposite of hungry is what word? Answer: Not hungry) were rejected to ensure participants performed the task by generating a lexical item rather than repeating the CI. Aside from correct answers, any of the following were noted:

#### Semantic paraphasia

1) an incorrect answer (i.e., giving a synonym when the question required an antonym or giving an answer unrelated semantically to the CI) to a question which was answered correctly at least once or 2) answers related semantically to the CI and/or correct response were considered errors if the same correct answer at least twice during the experiment. In both cases, an answer would be considered a semantic error if the participant spontaneously corrected their answer prior to completing the next trial, except when a participant offered additional correct responses.

#### Phonological paraphasia

a neologism, or an unattested English word consisting of segments that are properly articulated at a gross level.

#### Anomia

questions that the participant failed to answer or could not respond for any reason, except for trials where the participant either reported not being able to hear due to background noise or appeared to fall asleep, which were excluded from further analysis. The stimulation site in 494L (Extended Data Fig. 3c) where perceptual dysfunction was reported was excluded from all analyses of anomia.

#### Hesitations

trials which the participant produced a hesitation (e.g., “uh”, “um”) or dysfluency (e.g., stuttering, repetition or prolongation of the first segment) prior to responding. In addition, because all CI questions could be answered with a single word, any answers where this word was preceded by filler words (e.g., Question: The opposite of hungry is what word? Answer: “The opposite of hungry is full.”) were considered trials containing hesitations.

#### Qualitative results

any site where stimulation resulted in a motor effect (e.g., speech arrest, gaze deviation, dysarthria, etc.) as defined by a neurosurgeon (J.D.W.G.). Such effects were categorized as either ‘speech motor’ (i.e., speech arrest, orofacial tetanus, involuntary orofacial movement, sensation of orofacial movement, dysarthria) or ‘gaze motor’ (i.e., involuntary head and/or eye movement). Any site which resulted in a sensation or movement that indicated the delivery of DES to the participant was not included in behavioral analyses. Descriptions of all qualitative motor effects are provided in Extended Data Table 2.

Finally, any site displaying a higher rate of semantic paraphasia, anomia, hesitations, and/or neologisms in stimulated versus control trials (minimum increase: 1%) was considered to display an increase in errors.

#### Behavioral Analysis

Inter-turn gap duration was defined as the duration between question offset and answer onset (excluding any hesitations) and was computed for all trials where an answer was produced (i.e., trials without anomia). Stimulus-response latency, defined as the duration between stimulation offset and answer onset, was calculated for all stimulated trials.

Stimulation-induced differences in gap duration at each site were assessed in early and late CI trials separately, as CI position significantly affects gap timing^7,10,11^. Statistical significance in gap duration between control and stimulated trials was determined using a three-step process 1) rejecting outliers in the gap duration distributions for control and stimulated trials separately (i.e., any gaps greater than 2 interquartile ranges above the 75^th^ percentile or less than 2 interquartile ranges below the 25^th^ percentile), 2) identifying sites where the median gap duration in control and stimulated trials differed by at least 100 ms, and 3) performing rank-sum tests (α = 0.05) on the outlier-rejected distributions of gap duration in control and stimulated trials. Behavioral analyses were not performed unless there were at least 8 answered stimulated and 8 answered control trials after outlier rejection. Due to restricted study times, we presented participants with more early trials than late trials (at an ∼2:1 ratio); consequently, many sites did not have sufficient late trials for behavioral analysis. Trial numbers, gap duration summary statistics, and other behavioral data for each stimulation site is presented in Extended Data Table 2.

## Statistical Analyses

All analyses, including permutation tests (100,000 iterations), were performed in MATLAB (Mathworks, Natick, MA) using custom scripts. Rank-sum tests were performed with the ‘ranksum’ function, correlation analyses were performed with the ‘corr’ function, and logistic regression was performed using generalized linear modelling (‘fitglm’ function; binomial distribution, logit linkage function, and maximum of 1000 iterations specified). For all logistic regression analyses, the occurrence of stimulation-induced increases in gap duration was used as the categorical response variable and predictor variables were defined as follows:

1) Stimulation intensity vs. increased gap duration: Stimulation voltage was used as a single continuous predictor variable.
2) Clinical condition vs. increased gap duration: Stimulation sites were divided into three groups according to whether participants were being treated for epilepsy, a brain tumor, or essential tremor/Parkinson’s disease (combined due to limited sample size) (Extended Data Table 1). These groups were then used as categorical dummy variables, with the combined essential tremor/Parkinson’s disease group designated as the reference category.
3) Stimulator type vs. increased gap duration: Stimulation sites were divided into three groups according to whether DES was delivered via ECoG electrodes, a handheld stimulator, or sEEG electrodes. These groups were then used as categorical dummy variables, with the sEEG group designated as the reference category.

## Non-Author Contributions

We thank members of the Long laboratory as well as Taylor Abel, Frank Guenther, Elnaz Hozhabri, Jelena Krivokapić, Kenway Louie, Felix Moll, Caroline Niziolek, and Shy Shoham for comments on earlier versions of this manuscript. We also thank Joel Berger, Haiming Chen, Phillip Gander, Christopher Garcia, Matthew Howard III, My Hieu Kien Huynh, Kenji Ibayashi, Zahra Jourahmad, Hiroto Kawasaki, Mac McKay, Kirill Nourski, Ariane Rhone, and Andrea Rohl for help with data collection.

## AUTHOR CONTRIBUTIONS

Conceptualization: GAC, MAL

Data curation: GAC, CKK, FT

Formal analysis: GAC, CKK, FT, DC

Funding acquisition: JDWG, MAL

Investigation: GAC, DC, JDWG

Methodology: GAC, CKK, FT, DC, MAL, JDWG

Project administration: GAC, JDWG

Resources: JDWG, MAL

Software: GAC, CKK, FT, DC

Supervision: JDWG, MAL

Validation: GAC, CKK, JDWG

Visualization: GAC, MAL

Writing – original draft: GAC

Writing – review & editing: GAC, CKK, FT, DC, JDWG, MAL

## DATA AVAILABILITY

Due to concerns about patient confidentiality, data are not publicly available; however, the corresponding author will provide deidentified behavioral data upon request.

## CODE AVAILABILITY

All MATLAB code used in this study will be provided upon request to the corresponding author.

