## Extended Data Table 1 for "A frontal cortical network is critical for language planning during spoken interaction"

**Extended Data Table 1.** Participant demographics and clinical information.

| Number | Age | Sex | Stimulator Type | Language Dominance (Method) | Handedness | Pathology/Diagnosis | Tumor Location/Seizure Loci | Notes |
|---|---|---|---|---|---|---|---|---|
| 460L | 51 | M | Chronic subdural ECoG | LH (Wada testing) | right | Epilepsy (gliosis) | Left mesial temporal lobe | |
| 494L | 30 | M | Acute Handheld | LH (Wada testing) | right (+70) | Hippocampal sclerosis (epilepsy) | Left hippocampus | |
| 524R | 18 | M | Chronic SEEG | LH (fMRI) | left (-30) | Epilepsy | Bilateral multifocal | |
| 561B | 19 | M | Chronic SEEG | LH (fMRI) | right | Epilepsy | Right parieto-occipital lobe | |
| 567R | 33 | M | Chronic SEEG | Unknown | right (+90) | Epilepsy | Right peri-sylvian (posterior frontal-anterior parietal medial operculum/posterior insula) | |
| 579B | 12 | F | Chronic SEEG | Unknown | left | Epilepsy | Right occipital | |
| 585L | 38 | F | Chronic subdural ECoG | LH (Wada testing) | right | Epilepsy | left middle hippocampus | |
| 625L | 22 | F | Chronic SEEG | Bihemispheric (Wada testing) | right | Epilepsy | left anterior medial temporal lobe | |
| 634L | 20 | F | Chronic SEEG | LH (Wada testing) | right | Epilepsy | left amygdala and hippocampus and insula | |
| 641L | 37 | M | Acute Handheld | Unknown | right (+70) | Astrocytoma | left frontal | Stimulation timing was estimated using from video of stimulator |
| 645L | 40 | F | Chronic SEEG | Bihemispheric (Wada testing) | mixed | Epilepsy; Hydrocephalus | bilateral amygdala | |
| 640L | 43 | F | Chronic SEEG | Bihemispheric (Wada testing) | right | Epilepsy | left amygdala | Patient reported occassional perception of scalp tingling during experiment, but did not appear to relate to stimulation delivery |
| 668L | 55 | F | Acute Handheld | Unknown | right | Glioblastoma Multiforme | left parietal | |
| 673R | 43 | M | Acute Handheld | Unknown | right | Glioblastoma Multiforme | right frontal (precentral gyrus) | |
| 671DBS | 71 | M | Acute subdural ECoG | Unknown | right | Parkinson's Disease | N/A | |
| 679L | 57 | F | Acute Handheld | Unknown | right | Esophageal CA metastasis; Hydrocephalus | Left frontoparietal | |
| 683DBS | 71 | F | Acute subdural ECoG | Unknown | right | Parkinson's Disease | N/A | |
| 684DBS | 54 | F | Acute subdural ECoG | Unknown | right | Parkinson's Disease | N/A | Vietnamese native speaker (English L2); CI task data not analyzed for stimulation effects in reaction time or errors and second stimulation site lacking gross motor deficits was not analyzed |
| 682DBS | 68 | M | Acute subdural ECoG | Unknown | right | Parkinson's Disease | N/A | |
| 672B | 34 | F | Chronic SEEG | LH (Wada testing) | right | Epilepsy | Right mesial temporal lobe | |
| 701L | 67 | M | Acute Handheld | Unknown | right | Lung CA Metastasis | Left temporal lobe | |
| 706DBS | 56 | F | Acute subdural ECoG | Unknown | right | Essential Tremor | N/A | |
| 716DBS | 51 | M | Acute subdural ECoG | Unknown; speech motor site on left precentral gyrus | left | Essential Tremor | N/A | |
