## Extended Data Table 2 for "A frontal cortical network is critical for language planning during spoken interaction"

**Extended Data Table 2**. Additional anatomical information and summary statistics for each stimulation site.

| Participant | Site | CIT168 Coregistered Coordinates | | | DKT Label | Condition | Inter-Turn Gap Duration | | | | Semantic Paraphasia | | Anomia | | Hesitations | | Neologisms | | Result Summary |
|---|---|---|---|---|---|---|---|---|---|---|---|---|---|---|---|---|---|---|---|
| | | X | Y | Z | | | Early CI Trials (s) | p-value | Late CI Trials (s) | p-value | # | Rate | # | Rate | # | Rate | # | Rate | |
| 460 | 204, Contact 1 | -46.89 | 43.22 | 24.51 | left pars triangularis | Control | 0.374 | 0.0088 | 0.675 | 0.0095 | 0 | 0 | 0 | 0 | 2 | 0.044 | 0 | 0 | longer gap durations; increased errors |
| | 204, Contact 2 | -52.93 | 37.73 | 16.39 | left caudal middle frontal gyrus | Stimulated | 0.555 | | 1.635 | | 1 | 0.023 | 1 | 0.023 | 5 | 0.114 | 0 | 0 | |
| 494 | 1, Contact 1 | -57.26 | 25.85 | 3.86 | left pars triangularis | Control | 0.133 | 0.0010 | 0.373 | 0.0003 | 1 | 0.031 | N/A | N/A | 1 | 0.031 | 0 | 0 | longer gap durations; increased errors; perceptual deficits |
| | 1, Contact 2 | -57.69 | 19.43 | -1.98 | left pars triangularis | Stimulated | 0.454 | | 0.785 | | 2 | 0.063 | N/A | N/A | 3 | 0.094 | 0 | 0 | |
| | 2, Contact 1 | -56.66 | 25.91 | 7.55 | left pars triangularis | Control | 0.094 | 0.8197 | insufficient trials | | 0 | 0 | 0 | 0 | 0 | 0 | 0 | 0 | increased errors |
| | 2, Contact 2 | -55.12 | 14.94 | 4.41 | left pars opercularis | Stimulated | 0.078 | | | | 2 | 0.125 | 2 | 0.125 | 0 | 0 | 0 | 0 | |
| | 3, Contact 1 | -60.71 | 11.91 | 10.79 | left precentral gyrus | Control | N/A | N/A | N/A | N/A | N/A | N/A | N/A | N/A | N/A | N/A | N/A | N/A | motor deficit (speech arrest) |
| | 3, Contact 2 | -60.63 | 5.75 | 7.32 | left precentral gyrus | Stimulated | N/A | | N/A | | N/A | N/A | N/A | N/A | N/A | N/A | N/A | N/A | |
| | 4, Contact 1 | -62.36 | 2.01 | 12.31 | left precentral gyrus | Control | N/A | N/A | N/A | N/A | N/A | N/A | N/A | N/A | N/A | N/A | N/A | N/A | motor deficit (dysarthria) |
| | 4, Contact 2 | -66.41 | -7.89 | 8.65 | left precentral gyrus | Stimulated | N/A | | N/A | | N/A | N/A | N/A | N/A | N/A | N/A | N/A | N/A | |
| 524 | 118, Contact 1 | 35.96 | -22.06 | 17.88 | right insula | Control | 0.232 | 0.2675 | insufficient trials | | 0 | 0 | 0 | 0 | 3 | 0.059 | 0 | 0 | increased errors |
| | 118, Contact 2 | 40.88 | -20.78 | 20.23 | right postcentral gyrus (parietal operculum) | Stimulated | 0.214 | | | | 1 | 0.027 | 0 | 0 | 0 | 0 | 0 | 0 | |
| | 82, Contact 1 | 37.44 | 8.03 | 57.37 | right caudal middle frontal gyrus | Control | 0.195 | 5.17E-06 | 0.329 | 0.0058 | 0 | 0 | 0 | 0 | 1 | 0.019 | 0 | 0 | longer gap durations; increased errors |
| | 82, Contact 2 | 42.43 | 9.88 | 59.37 | right caudal middle frontal gyrus | Stimulated | 0.840 | | 0.741 | | 0 | 0 | 2 | 0.039 | 3 | 0.059 | 0 | 0 | |
| 561 | 19, Contact 1 | 7.06 | -22.97 | 29.95 | right posterior cingulate gyrus | Control | 0.846 | 0.1152 | insufficient trials | | 0 | 0 | 0 | 0 | 2 | 0.059 | 0 | 0 | no effect |
| | 19, Contact 2 | 9.79 | -22.44 | 33.49 | right posterior cingulate gyrus (white matter) | Stimulated | 0.547 | | | | 0 | 0 | 0 | 0 | 1 | 0.028 | 0 | 0 | |
| | 216, Contact 1 | -0.36 | 0.43 | 31.63 | left posterior cingulate gyrus | Control | 0.818 | 6.16E-07 | insufficient trials | | 0 | 0 | 0 | 0 | 4 | 0.085 | 0 | 0 | longer gap durations; increased errors |
| | 216, Contact 2 | -4.35 | 1.67 | 33.78 | left posterior cingulate gyrus | Stimulated | 2.060 | | | | 0 | 0 | 0 | 0 | 9 | 0.205 | 0 | 0 | |
| | 225, Contact 1 | -30.93 | 11.95 | 50.83 | left caudal middle frontal gyrus | Control | 0.667 | 0.0156 | 1.126 | 0.9654 | 0 | 0 | 0 | 0 | 2 | 0.039 | 0 | 0 | longer gap durations; increased errors |
| | 225, Contact 2 | -35.77 | 10.24 | 52.16 | left caudal middle frontal gyrus | Stimulated | 0.874 | | 1.073 | | 0 | 0 | 0 | 0 | 9 | 0.188 | 0 | 0 | |
| | 81, Contact 1 | 57.77 | 33.36 | 5.07 | right pars triangularis | Control | 0.704 | 0.5169 | insufficient trials | | 0 | 0 | 0 | 0 | 0 | 0 | 0 | 0 | no effect |
| | 81, Contact 2 | 63.78 | 32.52 | 10.09 | right pars triangularis | Stimulated | 0.790 | | | | 0 | 0 | 0 | 0 | 0 | 0 | 0 | 0 | |
| 567 | 155, Contact 1 | 33.07 | 12.80 | 65.59 | right caudal middle frontal gyrus | Control | 0.922 | 0.1495 | 2.297 | 0.4807 | 0 | 0 | 2 | 0.041 | 11 | 0.224 | 0 | 0 | no effect |
| | 155, Contact 2 | 39.04 | 10.59 | 65.61 | right caudal middle frontal gyrus | Stimulated | 1.094 | | 1.398 | | 0 | 0 | 0 | 0 | 8 | 0.182 | 0 | 0 | |
| | 177, Contact 1 | 36.67 | 7.94 | 51.59 | right caudal middle frontal gyrus | Control | N/A | N/A | N/A | N/A | N/A | N/A | N/A | N/A | N/A | N/A | N/A | N/A | motor deficit (head and gaze movement) |
| | 177, Contact 2 | 41.72 | 7.48 | 54.76 | right caudal middle frontal gyrus | Stimulated | N/A | | N/A | | N/A | N/A | N/A | N/A | N/A | N/A | N/A | N/A | |
| 579 | 46, Contact 1 | -49.91 | -3.15 | 46.59 | left precentral gyrus | Control | N/A | N/A | N/A | N/A | N/A | N/A | N/A | N/A | N/A | N/A | N/A | N/A | motor deficit (head and gaze movement) |
| | 46, Contact 2 | -52.17 | -1.99 | 47.12 | left precentral gyrus | Stimulated | N/A | | N/A | | N/A | N/A | N/A | N/A | N/A | N/A | N/A | N/A | |
| | 94, Contact 1 | -48.95 | 33.60 | 3.88 | left pars triangularis | Control | 0.243 | 0.8644 | insufficient trials | | 0 | 0 | 0 | 0 | 0 | 0 | 0 | 0 | increased errors |
| | 94, Contact 2 | -52.24 | 35.60 | 6.22 | left pars triangularis | Stimulated | 0.272 | | | | 0 | 0 | 0 | 0 | 1 | 0.067 | 0 | 0 | |
| 585 | 152, Contact 1 | -15.69 | 72.19 | -8.47 | left frontal pole | Control | 0.267 | 0.5487 | insufficient trials | | 0 | 0 | 0 | 0 | 0 | 0 | 0 | 0 | increased errors |
| | 152, Contact 2 | -25.21 | 69.34 | -7.39 | left frontal pole | Stimulated | 0.288 | | | | 0 | 0 | 0 | 0 | 1 | 0.032 | 0 | 0 | |
| | 230, Contact 1 | -41.14 | 19.83 | 56.73 | left caudal middle frontal gyrus | Control | 0.303 | 0.2032 | insufficient trials | | 0 | 0 | 0 | 0 | 0 | 0 | 0 | 0 | increased errors |
| | 230, Contact 2 | -49.12 | 9.18 | 56.10 | left caudal middle frontal gyrus | Stimulated | 0.337 | | | | 0 | 0 | 0 | 0 | 2 | 0.049 | 0 | 0 | |
| | 236, Contact 1 | -50.16 | 32.65 | 33.25 | left caudal middle frontal gyrus | Control | 0.560 | 0.2474 | insufficient trials | | 0 | 0 | 0 | 0 | 2 | 0.069 | 0 | 0 | no effect |
| | 236, Contact 2 | -56.41 | 27.46 | 26.59 | left caudal middle frontal gyrus | Stimulated | 0.489 | | | | 0 | 0 | 0 | 0 | 2 | 0.063 | 0 | 0 | |
| | 250, Contact 1 | -57.67 | 32.62 | -4.80 | left pars orbitalis | Control | 0.514 | 0.6182 | insufficient trials | | 0 | 0 | 0 | 0 | 0 | 0 | 0 | 0 | no effect |
| | 250, Contact 2 | -59.95 | 27.06 | 5.76 | left pars triangularis | Stimulated | 0.503 | | | | 0 | 0 | 0 | 0 | 0 | 0 | 0 | 0 | |
| 625 | 10, Contact 1 | -48.70 | 34.63 | -5.41 | left pars orbitalis | Control | 0.957 | 0.0001 | 1.169 | 0.4636 | 1 | 0.030 | 3 | 0.091 | 1 | 0.030 | 0 | 0 | shorter gap durations; increased errors |
| | 10, Contact 2 | -53.41 | 35.01 | -4.12 | left pars orbitalis | Stimulated | 0.150 | | 1.050 | | 4 | 0.103 | 4 | 0.103 | 2 | 0.051 | 0 | 0 | |
| | 63, Contact 1 | -52.01 | 7.50 | 33.84 | left precentral gyrus | Control | N/A | N/A | N/A | N/A | N/A | N/A | N/A | N/A | N/A | N/A | N/A | N/A | motor deficit (orofacial movement which the participant described as "a string pulling" face/mouth) |
| | 63, Contact 2 | -57.65 | 7.33 | 35.08 | left precentral gyrus | Stimulated | N/A | | N/A | | N/A | N/A | N/A | N/A | N/A | N/A | N/A | N/A | |
| | 80, Contact 1 | -38.00 | 19.54 | 6.76 | left insula | Control | 0.416 | 0.5089 | 1.243 | 0.9546 | 0 | 0 | 5 | 0.172 | 4 | 0.138 | 0 | 0 | increased errors |
| | 80, Contact 2 | -39.56 | 23.26 | 11.24 | left pars opercularis (frontal operculum) | Stimulated | 0.593 | | 1.045 | | 4 | 0.138 | 5 | 0.172 | 3 | 0.103 | 0 | 0 | |
| 634 | 23, Contact 1 | -3.75 | 32.34 | 19.90 | left caudal anterior cingulate gyrus | Control | 0.191 | 0.2424 | 0.459 | 1.0000 | 0 | 0 | 0 | 0 | 0 | 0 | 0 | 0 | no effect |
| | 23, Contact 2 | -8.39 | 33.37 | 21.14 | left caudal anterior cingulate gyrus | Stimulated | 0.149 | | 0.474 | | 0 | 0 | 0 | 0 | 0 | 0 | 0 | 0 | |
| | 40, Contact 1 | -40.76 | 2.38 | 41.06 | left caudal middle frontal gyrus (precentral sulcus) | Control | 0.117 | 1.54E-07 | 0.439 | 0.0006 | 1 | 0.029 | 1 | 0.029 | 0 | 0 | 0 | 0 | longer gap durations; increased errors |
| | 40, Contact 2 | -46.15 | 2.85 | 41.36 | left caudal middle frontal gyrus (precentral sulcus) | Stimulated | 3.445 | | 2.663 | | 1 | 0.028 | 0 | 0 | 5 | 0.139 | 0 | 0 | |
| 640 | 24, Contact 1 | 55.02 | -31.30 | 1.00 | right superior temporal gyrus | Control | 0.271 | 0.8467 | 0.524 | 0.0289 | 0 | 0 | 0 | 0 | 0 | 0 | 0 | 0 | no effect |
| | 24, Contact 2 | 60.42 | -31.32 | 1.59 | right superior temporal gyrus | Stimulated | 0.263 | | 0.433 | | 0 | 0 | 0 | 0 | 0 | 0 | 0 | 0 | |
| | 39, Contact 1 | -49.70 | -0.59 | -21.49 | left superior temporal gyrus | Control | 0.255 | 0.8090 | 0.525 | 0.4025 | 0 | 0 | 0 | 0 | 0 | 0 | 0 | 0 | no effect |
| | 39, Contact 2 | -56.16 | -0.64 | -21.44 | left middle temporal gyrus | Stimulated | 0.280 | | 0.486 | | 0 | 0 | 0 | 0 | 1 | 0.028 | 0 | 0 | |
| | 74, Contact 1 | -47.93 | -28.91 | 9.84 | left transverse temporal gyrus | Control | 0.323 | 0.1424 | 0.535 | 0.7350 | 0 | 0 | 0 | 0 | 0 | 0 | 0 | 0 | no effect |
| | 74, Contact 2 | -53.74 | -28.25 | 11.52 | left transverse temporal gyrus | Stimulated | 0.349 | | 0.490 | | 0 | 0 | 0 | 0 | 0 | 0 | 0 | 0 | |
| | 90, Contact 1 | -45.57 | -29.35 | -19.48 | left fusiform gyrus | Control | 0.312 | 3.46E-07 | 0.563 | 0.0001 | 0 | 0 | 0 | 0 | 2 | 0.053 | 0 | 0 | longer gap durations; increased errors |
| | 90, Contact 2 | -50.95 | -29.92 | -20.13 | left inferior temporal gyrus | Stimulated | 2.275 | | 2.724 | | 3 | 0.094 | 1 | 0.031 | 16 | 0.500 | 0 | 0 | |
| | 9, Contact 1 | 54.13 | 0.73 | -19.31 | right middle temporal gyrus | Control | 0.237 | 0.5301 | 0.510 | 0.9310 | 0 | 0 | 0 | 0 | 0 | 0 | 0 | 0 | no effect |
| | 9, Contact 2 | 60.00 | 0.44 | -19.34 | right middle temporal gyrus | Stimulated | 0.221 | | 0.418 | | 0 | 0 | 0 | 0 | 0 | 0 | 0 | 0 | |
| 641 | 1, Contact 1 | -56.48 | 31.29 | 20.92 | left pars triangularis | Control | 0.346 | 0.0146 | 1.013 | 0.3811 | 0 | 0.000 | 11 | 0.134 | 3 | 0.037 | 0 | 0 | longer gap durations; increased errors |
| | 1, Contact 2 | -55.84 | 34.99 | 16.90 | left pars triangularis | Stimulated | 0.578 | | 1.143 | | 1 | 0.014 | 8 | 0.113 | 4 | 0.056 | 0 | 0 | |
| 645 | 38, Contact 1 | -47.84 | 10.89 | -28.31 | left superior temporal gyrus | Control | 1.009 | 0.2797 | 1.180 | 0.4303 | 0 | 0 | 0 | 0 | 1 | 0.030 | 0 | 0 | increased errors |
| | 38, Contact 2 | -53.25 | 11.10 | -29.38 | left superior temporal gyrus | Stimulated | 0.951 | | 1.167 | | 0 | 0 | 0 | 0 | 4 | 0.091 | 0 | 0 | |
| | 45, Contact 1 | -51.18 | -19.14 | -11.73 | left middle temporal gyrus | Control | 0.730 | 0.4958 | 2.263 | 0.0092 | 1 | 0.027 | 0 | 0 | 1 | 0.027 | 0 | 0 | shorter gap durations; increased errors |
| | 45, Contact 2 | -58.39 | -19.54 | -11.27 | left middle temporal gyrus | Stimulated | 0.900 | | 1.033 | | 1 | 0.029 | 1 | 0.029 | 2 | 0.057 | 0 | 0 | |
| | 65, Contact 1 | -41.52 | -25.30 | 9.83 | left transverse temporal gyrus | Control | 0.448 | 0.3185 | 0.740 | 0.5114 | 0 | 0 | 1 | 0.026 | 1 | 0.026 | 0 | 0 | no effect |
| | 65, Contact 2 | -47.53 | -24.95 | 10.27 | left transverse temporal gyrus | Stimulated | 0.465 | | 0.751 | | 0 | 0 | 0 | 0 | 0 | 0.000 | 0 | 0 | |
| | 69, Contact 1 | -68.04 | -30.13 | 15.38 | left superior temporal gyrus | Control | 1.212 | 0.0389 | 1.704 | 0.1260 | 0 | 0 | 0 | 0 | 4 | 0.114 | 0 | 0 | shorter gap durations; increased errors |
| | 69, Contact 2 | -71.20 | -31.88 | 15.50 | left superior temporal gyrus | Stimulated | 0.539 | | 1.231 | | 0 | 0 | 1 | 0.028 | 1 | 0.028 | 0 | 0 | |
| 668 | 1, Contact 1 | -33.85 | -60.14 | 63.71 | left superior parietal cortex | Control | 0.100 | 0.1564 | insufficient trials | | 0 | 0 | 0 | 0 | 0 | 0 | 0 | 0 | no effect |
| | 1, Contact 2 | -34.04 | -63.45 | 61.21 | left superior parietal cortex | Stimulated | 0.110 | | | | 0 | 0 | 0 | 0 | 0 | 0 | 0 | 0 | |
| | 2, Contact 1 | -48.64 | -44.08 | 61.29 | left superior parietal cortex | Control | 0.092 | 0.1433 | insufficient trials | | 0 | 0 | 0 | 0 | 0 | 0 | 0 | 0 | no effect |
| | 2, Contact 2 | -45.57 | -44.74 | 65.00 | left superior parietal cortex | Stimulated | 0.060 | | | | 0 | 0 | 0 | 0 | 0 | 0 | 0 | 0 | |
| 671 | 1, Contact 1 | -60.51 | 20.10 | 7.58 | left pars triangularis | Control | 1.125 | 0.1285 | insufficient trials | | 3 | 0.143 | 4 | 0.190 | 5 | 0.238 | 0 | 0 | increased errors |
| | 1, Contact 2 | -59.33 | 22.19 | 15.29 | left pars triangularis | Stimulated | 0.788 | | | | 2 | 0.100 | 3 | 0.150 | 6 | 0.300 | 0 | 0 | |
| | 3, Contact 1 | -56.86 | 24.11 | 24.97 | left caudal middle frontal gyrus | Control | 1.002 | 0.0191 | insufficient trials | | 3 | 0.120 | 4 | 0.160 | 9 | 0.360 | 0 | 0 | longer gap durations; increased errors |
| | 3, Contact 2 | -53.65 | 26.31 | 32.11 | left caudal middle frontal gyrus | Stimulated | 2.932 | | | | 5 | 0.250 | 6 | 0.300 | 9 | 0.450 | 0 | 0 | |
| 672 | 136, Contact 1 | -49.18 | 3.16 | -23.47 | left superior temporal gyrus | Control | 0.266 | 0.2798 | 0.981 | 0.9039 | 1 | 0.0278 | 4 | 0.111 | 0 | 0 | 0 | 0 | no effect |
| | 136, Contact 2 | -54.09 | 3.26 | -24.04 | left middle temporal gyrus | Stimulated | 0.565 | | 0.993 | | 1 | 0.0294 | 1 | 0.029 | 0 | 0 | 0 | 0 | |
| | 151, Contact 1 | -64.12 | -18.03 | -9.49 | left middle temporal gyrus | Control | 0.568 | 0.3233 | 0.727 | 0.5181 | 0 | 0 | 1 | 0.027 | 1 | 0.027 | 0 | 0 | no effect |
| | 151, Contact 2 | -68.60 | -18.29 | -8.29 | left middle temporal gyrus | Stimulated | 0.369 | | 0.867 | | 0 | 0 | 0 | 0 | 0 | 0 | 0 | 0 | |
| | 160, Contact 1 | -59.34 | -49.14 | 1.41 | left middle temporal gyrus | Control | 0.373 | 0.2798 | 0.702 | 0.2970 | 0 | 0 | 1 | 0.026 | 0 | 0 | 0 | 0 | no effect |
| | 160, Contact 2 | -64.28 | -50.10 | 2.47 | left middle temporal gyrus | Stimulated | 0.658 | | 0.978 | | 0 | 0 | 1 | 0.028 | 0 | 0 | 0 | 0 | |
| 673 | 1, Contact 1 | 45.64 | -17.54 | 66.52 | right precentral gyrus | Control | 1.045 | 0.9440 | insufficient trials | | 0 | 0 | 0 | 0 | 0 | 0 | 0 | 0 | increased errors |
| | 1, Contact 2 | 41.58 | -15.82 | 67.58 | right precentral gyrus | Stimulated | 1.035 | | | | 0 | 0 | 1 | 0.045 | 0 | 0 | 0 | 0 | |
| | 2, Contact 1 | 28.80 | -10.88 | 74.55 | right precentral gyrus | Control | 1.115 | 0.5297 | insufficient trials | | 0 | 0 | 0 | 0 | 0 | 0 | 0 | 0 | no effect |
| | 2, Contact 2 | 33.35 | -11.87 | 73.08 | right precentral gyrus | Stimulated | 1.062 | | | | 0 | 0 | 0 | 0 | 0 | 0 | 0 | 0 | |
| 679 | 1, Contact 1 | -39.94 | -21.80 | 67.02 | left precentral gyrus | Control | 0.110 | 0.2158 | insufficient trials | | 0 | 0 | 1 | 0.050 | 0 | 0 | 0 | 0 | increased errors |
| | 1, Contact 2 | -42.51 | -20.31 | 63.58 | left precentral gyrus | Stimulated | -0.145 | | | | 1 | 0.048 | 0 | 0 | 1 | 0.048 | 0 | 0 | |
| | 2, Contact 1 | -51.88 | -11.48 | 51.06 | left precentral gyrus | Control | -0.022 | 0.9342 | insufficient trials | | 0 | 0 | 2 | 0.087 | 0 | 0 | 0 | 0 | increased errors |
| | 2, Contact 2 | -53.99 | -10.80 | 47.81 | left precentral gyrus | Stimulated | -0.009 | | | | 0 | 0 | 0 | 0 | 2 | 0.100 | 0 | 0 | |
| 682 | 1, Contact 1 | -66.65 | 1.77 | 12.08 | left precentral gyrus | Control | 0.958 | 0.0018 | insufficient trials | | 5 | 0.152 | 2 | 0.061 | 4 | 0.121 | 1 | 0.030 | shorter gap durations |
| | 1, Contact 2 | -65.44 | 6.50 | 19.54 | left precentral gyrus | Stimulated | 0.276 | | | | 3 | 0.097 | 2 | 0.065 | 3 | 0.097 | 0 | 0 | |
| | 3, Contact 1 | -62.29 | 10.66 | 28.29 | left precentral gyrus | Control | 0.508 | 0.7420 | 0.765 | 0.9015 | 1 | 0.031 | 3 | 0.094 | 4 | 0.125 | 0 | 0 | increased errors |
| | 3, Contact 2 | -60.64 | 12.85 | 35.83 | left precentral gyrus | Stimulated | 0.520 | | 0.881 | | 3 | 0.088 | 2 | 0.059 | 5 | 0.147 | 0 | 0 | |
| | 5, Contact 1 | -56.77 | 16.54 | 41.96 | left caudal middle frontal gyrus | Control | 0.347 | 0.2607 | insufficient trials | | 0 | 0 | 1 | 0.030 | 2 | 0.061 | 0 | 0 | increased errors |
| | 5, Contact 2 | -51.36 | 18.41 | 46.54 | left caudal middle frontal gyrus | Stimulated | 0.707 | | | | 0 | 0 | 2 | 0.067 | 2 | 0.067 | 0 | 0 | |
| 683 | 1, Contact 1 | -68.20 | -24.11 | 38.50 | left postcentral gyrus | Control | 0.141 | 0.8245 | insufficient trials | | 3 | 0.091 | 4 | 0.121 | 4 | 0.121 | 0 | 0 | increased errors |
| | 1, Contact 2 | -63.91 | -15.04 | 43.70 | left postcentral gyrus (central sulcus) | Stimulated | 0.042 | | | | 1 | 0.032 | 4 | 0.129 | 3 | 0.097 | 2 | 0.065 | |
| 684 | 1, Contact 1 | -68.04 | -25.72 | 33.31 | left inferior parietal cortex | Control | N/A | N/A | N/A | N/A | N/A | N/A | N/A | N/A | N/A | N/A | N/A | N/A | motor deficit (involuntary vocalization) |
| | 1, Contact 2 | -65.05 | -17.86 | 39.50 | left postcentral gyrus | Stimulated | N/A | | N/A | | N/A | N/A | N/A | N/A | N/A | N/A | N/A | N/A | |
| 701 | 1, Contact 1 | -67.47 | -50.31 | -2.13 | left middle temporal gyrus | Control | 0.066 | 0.0489 | insufficient trials | | 0 | 0 | 0 | 0 | 0 | 0 | 0 | 0 | no effect |
| | 1, Contact 2 | -68.49 | -47.24 | -2.97 | left middle temporal gyrus | Stimulated | 0.012 | | | | 0 | 0 | 0 | 0 | 0 | 0 | 0 | 0 | |
| | 2, Contact 1 | -66.80 | -57.81 | 7.93 | left middle temporal gyrus | Control | -0.043 | 0.0477 | insufficient trials | | 0 | 0 | 0 | 0 | 0 | 0 | 0 | 0 | no effect |
| | 2, Contact 2 | -67.72 | -54.07 | 8.28 | left middle temporal gyrus | Stimulated | -0.015 | | | | 0 | 0 | 0 | 0 | 0 | 0 | 0 | 0 | |
| 706 | 1, Contact 1 | -69.23 | -10.86 | -2.88 | left superior temporal gyrus | Control | N/A | N/A | N/A | N/A | N/A | N/A | N/A | N/A | N/A | N/A | N/A | N/A | motor deficit (dysarthria) |
| | 1, Contact 2 | -65.64 | -2.41 | 5.50 | left precentral gyrus (lateral sulcus) | Stimulated | N/A | | N/A | | N/A | N/A | N/A | N/A | N/A | N/A | N/A | N/A | |
| | 3, Contact 1 | -64.64 | 5.74 | 14.36 | left precentral gyrus | Control | 0.212 | 0.0591 | 0.865 | 0.9682 | 0 | 0 | 0 | 0 | 1 | 0.033 | 0 | 0 | no effect |
| | 3, Contact 2 | -62.80 | 11.22 | 21.40 | left precentral gyrus | Stimulated | 0.168 | | 0.981 | | 0 | 0 | 0 | 0 | 1 | 0.034 | 0 | 0 | |
| | 5, Contact 1 | -56.87 | 16.60 | 31.52 | left pars opercularis | Control | 0.968 | 0.3865 | insufficient trials | | 0 | 0 | 0 | 0 | 2 | 0.125 | 0 | 0 | increased errors |
| | 5, Contact 2 | -53.30 | 24.38 | 41.35 | left caudal middle frontal gyrus | Stimulated | 0.365 | | | | 0 | 0 | 0 | 0 | 2 | 0.143 | 0 | 0 | |
| 716 | 1, Contact 1 | -58.10 | 35.85 | 0.56 | left pars triangularis | Control | 0.226 | 0.3348 | 0.651 | 0.3638 | 0 | 0 | 0 | 0.000 | 0 | 0 | 0 | 0 | increased errors |
| | 1, Contact 2 | -57.40 | 35.93 | 13.43 | left pars triangularis | Stimulated | 0.226 | | 0.501 | | 0 | 0 | 2 | 0.069 | 1 | 0.0345 | 0 | 0 | |
| | 3, Contact 1 | -54.37 | 37.11 | 23.28 | left caudal middle frontal gyrus | Control | 0.203 | 0.4034 | 0.661 | 0.7337 | 0 | 0 | 1 | 0.033 | 0 | 0 | 0 | 0 | no effect |
| | 3, Contact 2 | -49.89 | 37.77 | 30.17 | left caudal middle frontal gyrus | Stimulated | 0.309 | | 0.647 | | 0 | 0 | 1 | 0.033 | 0 | 0 | 0 | 0 | |
| | 5, Contact 1 | -46.60 | 38.48 | 37.23 | left caudal middle frontal gyrus | Control | 0.260 | 0.2892 | 0.809 | 0.2241 | 0 | 0 | 1 | 0.031 | 0 | 0 | 0 | 0 | no effect |
| | 5, Contact 2 | -39.15 | 37.22 | 46.35 | left caudal middle frontal gyrus | Stimulated | 0.318 | | 1.021 | | 0 | 0 | 0 | 0 | 0 | 0 | 0 | 0 | |
