## Extended Data Figure 1 for "A frontal cortical network is critical for language planning during spoken interaction"

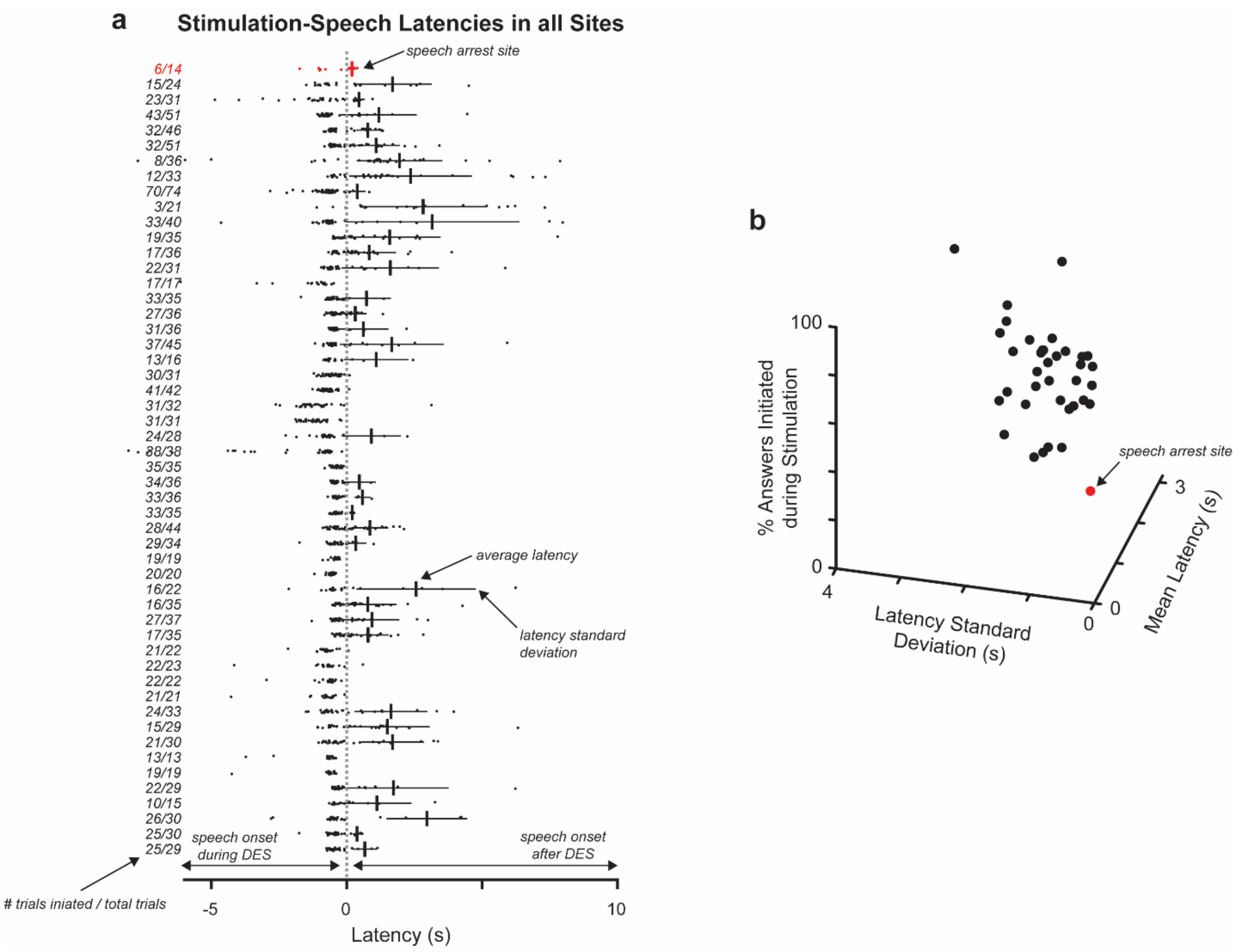


**Extended Data Figure 1. Additional quantification of stimulation-induced speech arrest.** (**a**) For all stimulation sites, mean stimulation-speech latency ± 1 standard deviation for trials where speech onset occurred after stimulation offset, with the proportion of trials answered while stimulation was ongoing at left; data for each site is presented in rows with dots representing latencies for individual trials. (**b**) Scatterplot reporting stimulation-speech latency summary statistics for each site.
