## Extended Data Figure 2 for "A frontal cortical network is critical for language planning during spoken interaction"

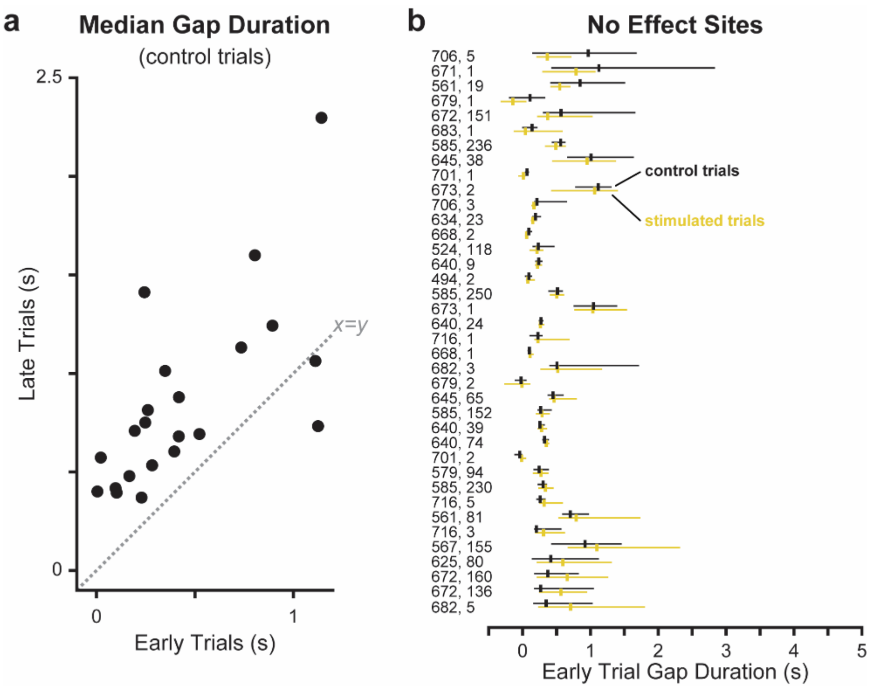


**Extended Data Figure 2. Additional quantification of inter-turn gap duration.** (**a**) For each participant, the median gap duration for all control trials pooled across stimulation sites when the CI was presented either question-medially (early) or question-finally (late); the median gap duration in early trials was significantly shorter than late trials across participants (p = 0.0002, sign-rank test). (**b**) Median gap duration and interquartile range for all sites where stimulation did not significantly affect this metric; participant and site number indicated in leftmost column (i.e., participant #, site #).
