## Extended Data Figure 3 for "A frontal cortical network is critical for language planning during spoken interaction"

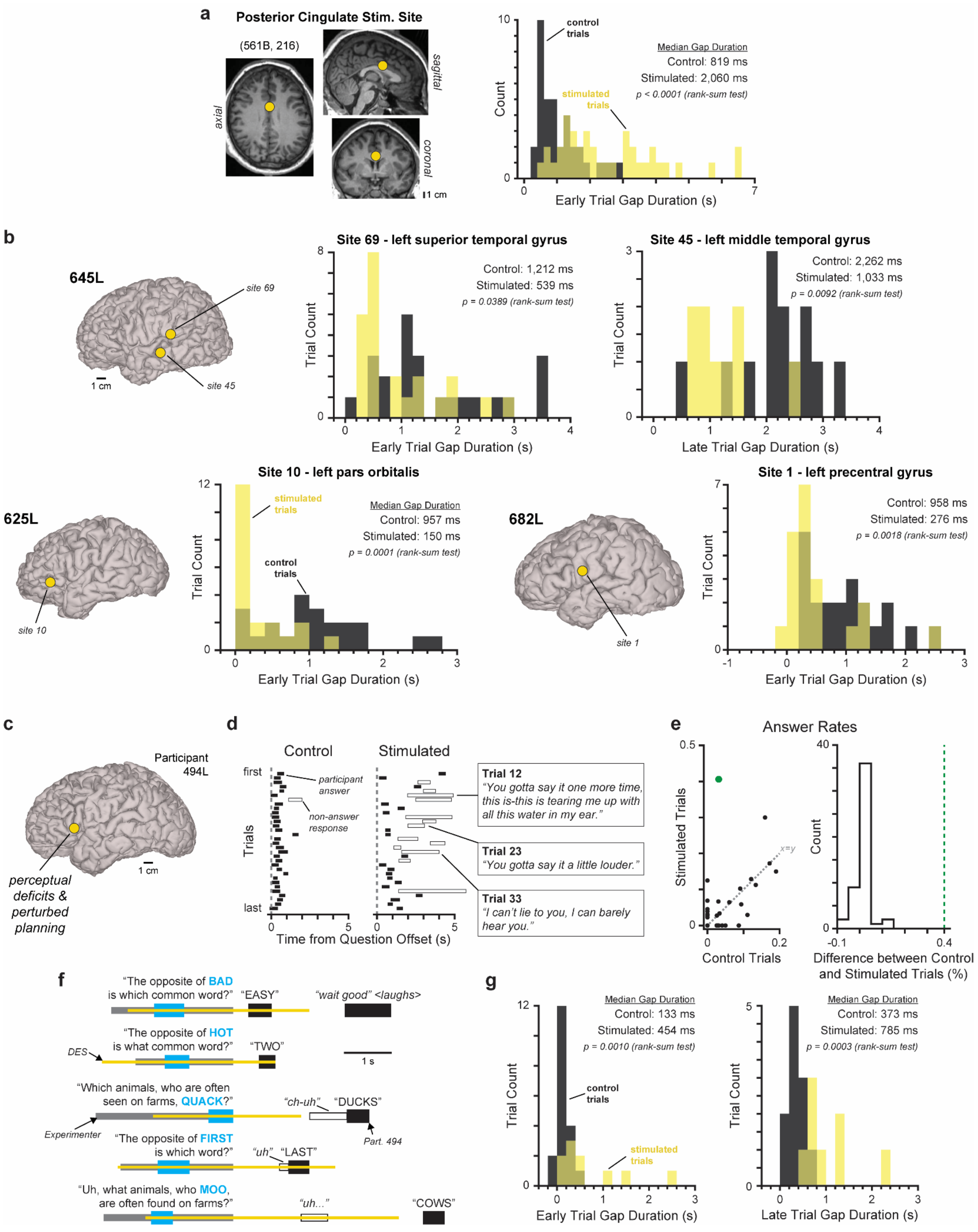

a
Posterior Cingulate Stim. Site
(561B, 216)
axial
sagittal
coronal
1 cm
control trials
stimulated trials
Median Gap Duration
Control: 819 ms
Stimulated: 2,060 ms
p < 0.0001 (rank-sum test)
Count
Early Trial Gap Duration (s)
b
645L
site 69
site 45
1 cm
Site 69 - left superior temporal gyrus
Control: 1,212 ms
Stimulated: 539 ms
p = 0.0389 (rank-sum test)
Trial Count
Early Trial Gap Duration (s)
Site 45 - left middle temporal gyrus
Control: 2,262 ms
Stimulated: 1,033 ms
p = 0.0092 (rank-sum test)
Trial Count
Late Trial Gap Duration (s)
625L
site 10
Site 10 - left pars orbitalis
stimulated trials
control trials
Median Gap Duration
Control: 957 ms
Stimulated: 150 ms
p = 0.0001 (rank-sum test)
Trial Count
Early Trial Gap Duration (s)
682L
site 1
Site 1 - left precentral gyrus
Control: 958 ms
Stimulated: 276 ms
p = 0.0018 (rank-sum test)
Trial Count
Early Trial Gap Duration (s)
c
Participant 494L
1 cm
perceptual deficits & perturbed planning
d
Control
Stimulated
first
last
Trials
participant answer
non-answer response
Time from Question Offset (s)
Trial 12
"You gotta say it one more time, this is-this is tearing me up with all this water in my ear."
Trial 23
"You gotta say it a little louder."
Trial 33
"I can't lie to you, I can barely hear you."
e
Answer Rates
Stimulated Trials
Control Trials
x=y
Count
Difference between Control and Stimulated Trials (%)
f
"The opposite of BAD is which common word?"
"EASY"
"wait good" <laughs>
"The opposite of HOT is what common word?"
"TWO"
DES
1 s
"Which animals, who are often seen on farms, QUACK?"
"ch-uh"
"DUCKS"
Experimenter
Part. 494
"The opposite of FIRST is which word?"
"uh"
"LAST"
"Uh, what animals, who MOO, are often found on farms?"
"uh..."
"COWS"
g
Median Gap Duration
Control: 133 ms
Stimulated: 454 ms
p = 0.0010 (rank-sum test)
control trials
stimulated trials
Trial Count
Early Trial Gap Duration (s)
Median Gap Duration
Control: 373 ms
Stimulated: 785 ms
p = 0.0003 (rank-sum test)
Trial Count
Late Trial Gap Duration (s)

**Extended Data Figure 3. Additional analysis of sites where stimulation altered inter-turn gap duration.** (**a**) Preoperative magnetic resonance images depicting the stimulation site within posterior cingulate gyrus (right) and a histogram of gap duration in control trials and trials when this site was stimulated (left). (**b**) Cortical surfaces from 3 participants depicting the location of sites where stimulation resulted in significantly shorter inter-turn gaps with histograms of gap duration in control and stimulated trials for each site. (**c**) Left lateral cortical surface of participant 494L with stimulation site indicated. (**d**) Answer timing in control trials (left) and trials when the site in (c) was stimulated (middle); example non-answer responses where the participant reported stimulation-induced perceptual dysfunction (right). (**e**) Proportion of control and stimulated trials answered (left) and the difference in answer rates (right) across participants; value related to site in (c) is indicated by green point and green dashed line. (**f**) Semantic paraphasia and hesitations resulting from stimulation of the site in (c). (**e**) Histograms of gap duration in control trials and stimulated trials for the site in (c).
